# Neuronal SNAP-23 scales hippocampal synaptic plasticity and memory

**DOI:** 10.1101/2022.08.04.502541

**Authors:** Mengjia Huang, Na-Ryum Bin, Jayant Rai, Ke Ma, Chun Hin Chow, Sarah Eide, Hidekiyo Harada, Jianbing Xiao, Daorong Feng, Hong-Shuo Sun, Zhong-Ping Feng, Herbert Y. Gaisano, Jeffrey E. Pessin, Philippe P. Monnier, Kenichi Okamoto, Liang Zhang, Shuzo Sugita

## Abstract

Soluble NSF Attachment protein REceptor (SNARE)-mediated membrane fusion plays a crucial role not only in presynaptic vesicle exocytosis but also in postsynaptic receptor delivery. The latter is considered particularly important for long-term synaptic plasticity and learning and memory, yet underlying mechanisms including the identity of the key SNARE proteins remain elusive. Here, we investigate the role of neuronal Synaptosomal-Associated Protein-23 (SNAP-23) by analyzing pyramidal-neuron specific SNAP-23 conditional knockout (cKO) mice. SNAP-23 immunostaining in postsynaptic spines was effectively decreased in the SNAP-23 cKO hippocampus. Electrophysiological analysis of SNAP-23 deficient neurons using acute hippocampal slices showed normal basal neurotransmission in CA3-CA1 synapses with unchanged AMPA and NMDA currents. Nevertheless, we found theta-burst stimulation induced long-term potentiation (LTP) was vastly diminished in SNAP-23 cKO. Moreover, unlike syntaxin-4 cKO mice in which both basal neurotransmission and LTP decrease manifested changes in a broad set of behavioral tasks, deficits of SNAP-23 cKO is more limited to spatial memory. Our data reveal that neuronal SNAP-23 is selectively crucial for synaptic plasticity and spatial memory without affecting basal glutamate receptor function.

## Introduction

Neuronal communication is essential for the function of the brain. Synaptic communications between neurons are dependent on a cellular process called exocytosis, wherein cargo - such as neurotransmitters stored in presynaptic vesicles - is delivered to the extracellular environment. Glutamate is the main excitatory neurotransmitter in the mammalian central nervous system. Presynaptically released glutamate binds to postsynaptic ionotropic or metabotropic receptors, which then exert their downstream effects to elicit activation of postsynaptic neurons.

Presynaptic vesicle exocytosis depends on the SNARE complex, a ternary protein complex, in which vesicular R-SNARE synaptobrevin/VAMP binds to target Q-SNAREs syntaxin-1 and SNAP-25 into a complex that drives membrane fusion resulting in rapid neurotransmitter releases. Recent evidence suggests that vesicular exocytosis is also involved in the regulation of postsynaptic glutamate receptors, α-amino-3-hydroxy-5-methyl-4-isoxazolepropionic acid receptors (AMPARs) and *N*-Methyl-D-aspartic acid receptors (NMDARs) (1). These receptors are known to undergo rounds of receptor recycling, being inserted or removed from the postsynaptic membrane thereby controlling synaptic strength and plasticity as required to maintain proper neural communications and functions (2–9).

Long-term potentiation (LTP) is the most widely studied form of synaptic plasticity and is believed to be the local circuitry correlate of learning and memory. LTP induction by high-frequency or theta-burst stimulation are known to change postsynaptic receptor numbers and composition (10). LTP results in NMDAR activation which further triggers AMPAR insertion into the membrane of postsynaptic neurons via exocytosis, which is thought to be also dependent on the SNARE complex. Moreover, there are evidence suggesting that the exocytosis of ionotropic glutamate receptors to the postsynaptic side is critical for the maintenance phase of LTP (11–13). Similarly, the endocytosis of ionotropic glutamate receptors on the postsynaptic side is critical for long-term depression (LTD) (14, 15). In addition to controlling the number of surface postsynaptic receptors, changes in the stoichiometry of postsynaptic neurotransmitter receptors have also been observed (15). Moreover, it has been shown that not only do AMPA receptor numbers change during events of synaptic plasticity (11–14), but also NMDA receptor subunit changes can occur, with evidence suggesting that the delivery rate varies among the receptor subunits (12, 16).

SNARE proteins are hypothesized to be involved in receptor delivery by mediating vesicle fusion and trafficking at the postsynaptic membrane (11, 16, 17). Nonetheless, the precise identities of SNARE proteins implicated in this process remain unclear. The synaptosomal-associated protein (SNAP) protein is a t-SNARE protein involved in vesicle fusion. There are 4 SNAP isoforms, SNAP-23, SNAP-25, SNAP 29, and SNAP-47, with nomenclature based on their molecular weight. SNAP-25 is the neuronal isoform (18) while SNAP-23, 29 and 47 are ubiquitous isoforms (19, 20). Lethality of a global SNAP-23 knockout and other isoforms in mice has led to approaches such as using shRNA-mediated acute knockdown or heterozygous mice to be employed for assessment of the function. As such, the role of SNAP-23 in the regulation of postsynaptic AMPA and NMDA currents has been particularly controversial in the field. For example, some argue for a role of SNAP-23 in NMDAR trafficking (21), while others counter that SNAP-25 plays this role (22). More importantly, the lack of *in vivo* animal model precludes any investigation of behavioral consequences of the SNAP-23 deletion.

In this study, we use CaMKIIα-Cre to generate a pyramidal neuron specific conditional SNAP-23 KO (cKO) mouse (23–25). Using this cKO mouse line, we analyze the potential postsynaptic function of SNAP-23 *in vitro* and *in vivo* using a combinatorial histological, electrophysiological and behavioral approaches. We show that SNAP-23 cKO results in defects of synaptic plasticity without impairing the basal transmission. We also show the correlation of synaptic deficits and the behavioral changes by comparing the phenotypes of SNAP-23 cKO mice with that of syntaxin-4 cKO mice which have reduced basal transmission as well as LTP (25). Our results demonstrate that the behavioral changes of SNAP-23 cKO is more selective than that of syntaxin-4 cKO.

## Results

### Generation of pyramidal neuron-specific SNAP-23 cKO mice

SNAP-23 null and germline SNAP-23 deletion mice result in pre-implantation embryonic lethality prior to embryonic day 3.5 (24, 26). Therefore, we took advantage of a conditional knockout system to circumvent the embryonic lethality of global knockout models in order to test the role of SNAP-23. To do this, a pyramidal neuron-specific knockout (KO) of SNAP-23 was generated by crossing SNAP-23 flox/flox mice in which exon 3 and 4 were flanked by loxP sites with a CaMKIIα-Cre mouse line. CaMKIIα-Cre is strongly expressed in pyramidal neurons, particularly in the CA1 region, of the hippocampus (23, 27). This pyramidal neuron-specific SNAP-23 conditional KO (cKO) line appeared visually normal at weaning and the mice were viable and fertile.

We attempted to confirm deletion of SNAP-23 protein in pyramidal neurons by immunohistochemistry of hippocampal sections using two-photon microscopy. SNAP-23 is expressed ubiquitously and was seen throughout the examined CA1 hippocampal region. Consistent with previous reports, SNAP-23 immunostaining showed colocalization with postsynaptic phalloidin staining (Figure 1A and 1B) (21), as well as appearing adjacent to phalloidin staining. Phalloidin is known to primarily label the periodic actin lattice in dendritic spines in acute hippocampal sections (28). CaMKIIα-Cre is expressed highly in the hippocampus (23, 29) as such, we consistently saw a decrease in SNAP-23 signals in the apical dendritic spines of CA1 neurons in the SNAP-23 cKO slices (Figure 1A and 1B). This decrease was also observed in the distal dendrites of the CA1 neurons (supplemental Figure 1). In addition, we saw a decrease in the SNAP-23 signal adjacent to phalloidin, suggesting that SNAP-23 deletion not only takes place in postsynaptic CA1 neurons but also in presynaptic neurons. Nevertheless, gross morphology of the entire hippocampus remained unchanged following SNAP-23 deletion as identified by cresyl violet staining [n = 17 for control, n = 15 for SNAP-23 cKO, independent t-test, area: t_(30)_ = 0.34, p = 0.735; density: t_(30)_ = 0.09, p = 0.929] (Figures 1C and 1D). Therefore, our SNAP-23 cKO line allows us to examine the functional role of neuronal SNAP-23 in the hippocampus in the absence of major structure abnormality.

**Figure 1.**
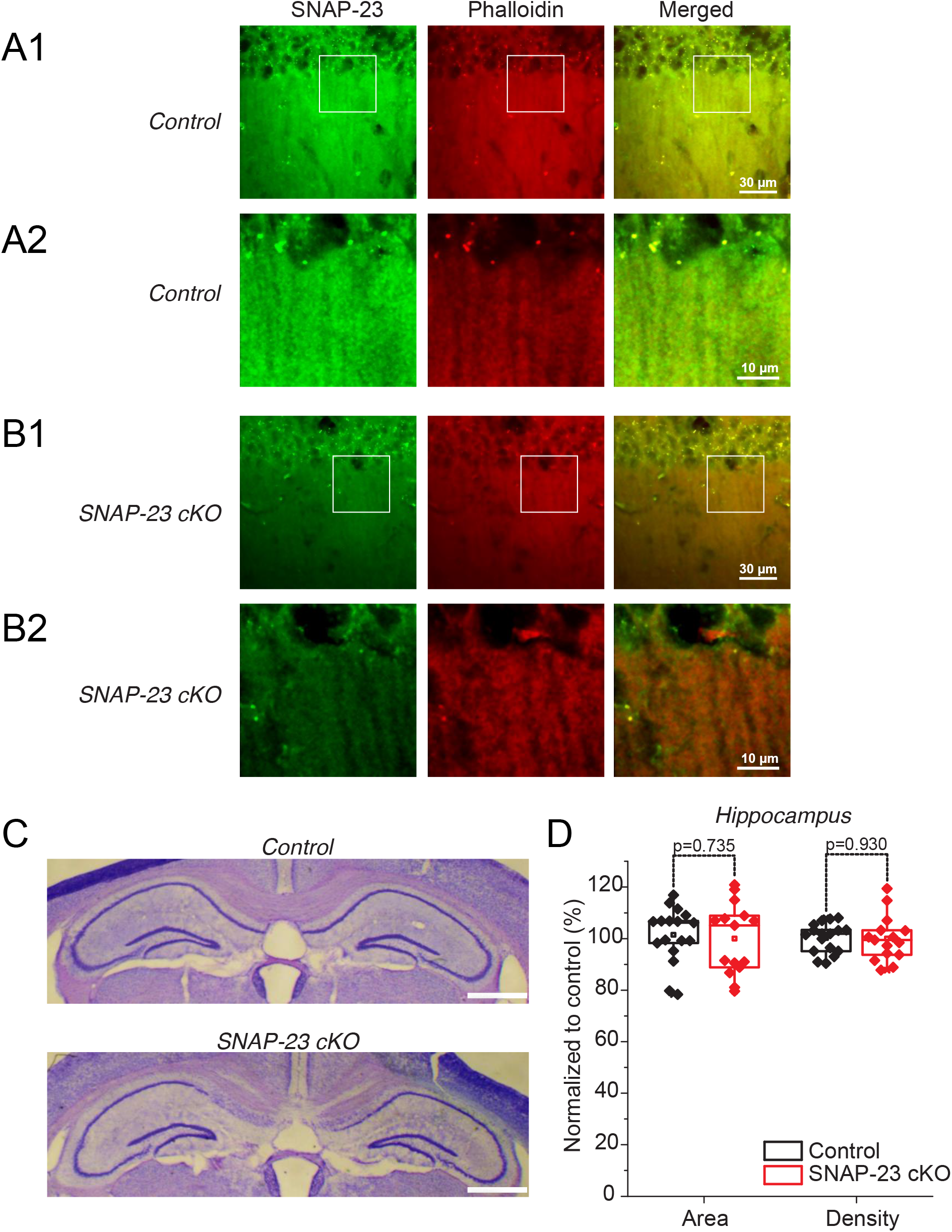
Generation of tissue-specific deletion of SNAP-23. SNAP-23 is specifically removed from CA1 cell dendrites. (A-B) Images obtained from coronal brain sections with the hippocampal CA1 region enlarged. SNAP-23 (green) and phalloidin (red) staining of CA1 apical dendrites from control (A) and SNAP-23 cKO (B) sections. Apical CA1 dendrite staining (A2, B2) shows a decrease in SNAP-23 cKO slice. (C) Coronal brain sections obtained from control and SNAP-23 cKO mice and stained with cresyl violet; bilateral hippocampal areas enlarged for illustration. Scale bar = 1 mm. (D) Area and density quantification of hippocampi of SNAP-23 cKO mice and control mice. Quantification was done in ImageJ (NIH, Bethesda, Maryland); hippocampi were manually selected, then area and intensity were quantified and normalized to control. n=17 for control, n=15 for cKO. No differences were observed between groups for area (p=0.735) or density (p=0.930); slices taken from 4 mice each. Error bars represent SEM.

### No significant change to evoked fEPSP in tissue-specific SNAP-23 cKO mice

We first tested if the CA3-CA1 synapses in SNAP-23 cKO mice exhibit changes in basal neurotransmission by using acute hippocampal slices with standard artificial cerebral spinal fluid (ACSF) perfusion. Field excitatory postsynaptic potentials (fEPSPs) were recorded from the apical dendrites of CA1 pyramidal neurons. A bipolar tungsten stimulating electrode was placed in the *striatum radiatum* of CA2 region to stimulate Schaffer collateral axons and paired stimuli of 50ms apart were delivered at incremental intensities from 10 to 150 μA (see Experimental Procedures). As more presynaptic axons are recruited with increasing stimulation intensities, we were able to quantify the amplitudes of the presynaptic fiber volley as a proxy of “input.” Input-output curves were generated to compare the fEPSP amplitudes and slopes between SNAP-23 control and cKO groups (Figures 2B and 2C). Increasing the amplitudes of the presynaptic fiber volley resulted in higher amplitudes and slopes of fEPSPs in the CA1 neurons, and such rise in both parameters was similarly observed in the control and the SNAP-23 cKO groups (Figures 2A–2C). In response to the paired stimuli, the ratios of the 2^nd^ vs. 1^st^ responses in the SNAP-23 cKO group was unchanged to that of the control group (Figures 2D and 2E). Together, these results indicate that SNAP-23 cKO pyramidal neurons result in no evident or significant changes to basal neurotransmission and short-term plasticity.

**Figure 2.**
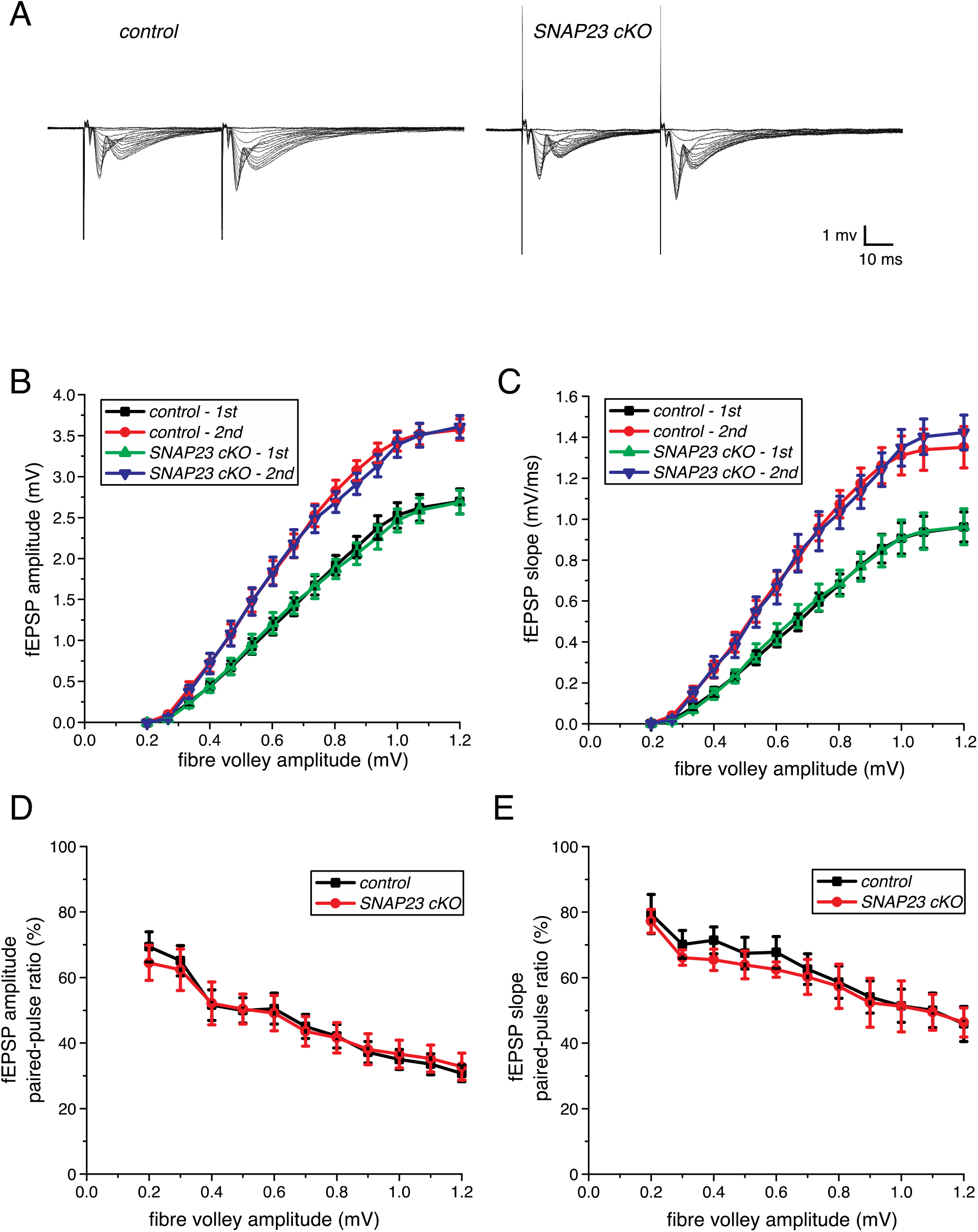
Tissue-specific deletion of SNAP-23 results in no significant changes to fEPSP measures of hippocampal CA1 pyramidal neurons. The Schaffer collateral axonal fibres were stimulated twice of 50ms apart and stimulating intensities varied successively from 10 to 150 μA. The resulting local fEPSP from CA1 pyramidal cell apical dendrites were recorded. (A) Sample traces from control and SNAP-23 cKO mice of dendritic fEPSP. (B) fEPSP amplitudes and (C) fEPSP slope were plotted against the presynaptic fibre volley amplitude. (E and F) Paired-pulse ratios of amplitudes (E) or slopes (F) were plotted against presynaptic fibre volley amplitudes. No significant changes between SNAP-23 cKO and controls were observed for all the measures. Error bars indicate SEM.

### Tissue-specific SNAP-23 deletion causes no change in both AMPAR and NMDAR-mediated fEPSPs or single cell EPSCs

A previous study using shRNA-mediated knockdown approach demonstrated that SNAP-23 is involved in the delivery of NMDARs to the postsynaptic membrane (21). Therefore, we investigated whether selective changes in AMPAR and/or NMDAR current could be observed using a pharmacological blockade approach in SNAP-23 cKO slices (Figure 3). A stable baseline recording was first obtained in standard ACSF, showing AMPAR-mediated fEPSPs predominantly. Switching the bath solution to Mg^2+^-free ACSF containing 100 μM picrotoxin allowed us to observe maximal ionotropic glutamate receptor-mediated fEPSPs by activating NMDAR-mediated responses and by blocking inhibitory GABAergic innervations. The evoked response revealed an enhanced glutamate receptor-mediated fEPSP with a characteristic prolonged epileptiform response containing multiple spikes in the early phase (Figure 3A). AMPARs largely mediate the multiple spikes in the early phase of the evoked response while the slow, long lasting later phase are largely mediated by NMDARs (25). Contributions of AMPAR and NMDAR-mediated fEPSPs were quantified by calculating the charge transfer, the integration of the area under the corresponding part of the traces (30). In contrast to previously reported results (21), we found no differences in AMPAR and NMDAR charge transfers in both the SNAP-23 cKO and the control groups (Figure 3C). The ratios of charge transfer from the AMPAR and the NMDAR-mediated responses were also not different between the control and the SNAP-23 cKO groups. Therefore, we found that SNAP-23 neither regulates basal AMPA or NMDA currents in postsynaptic neurons.

**Figure 3.**
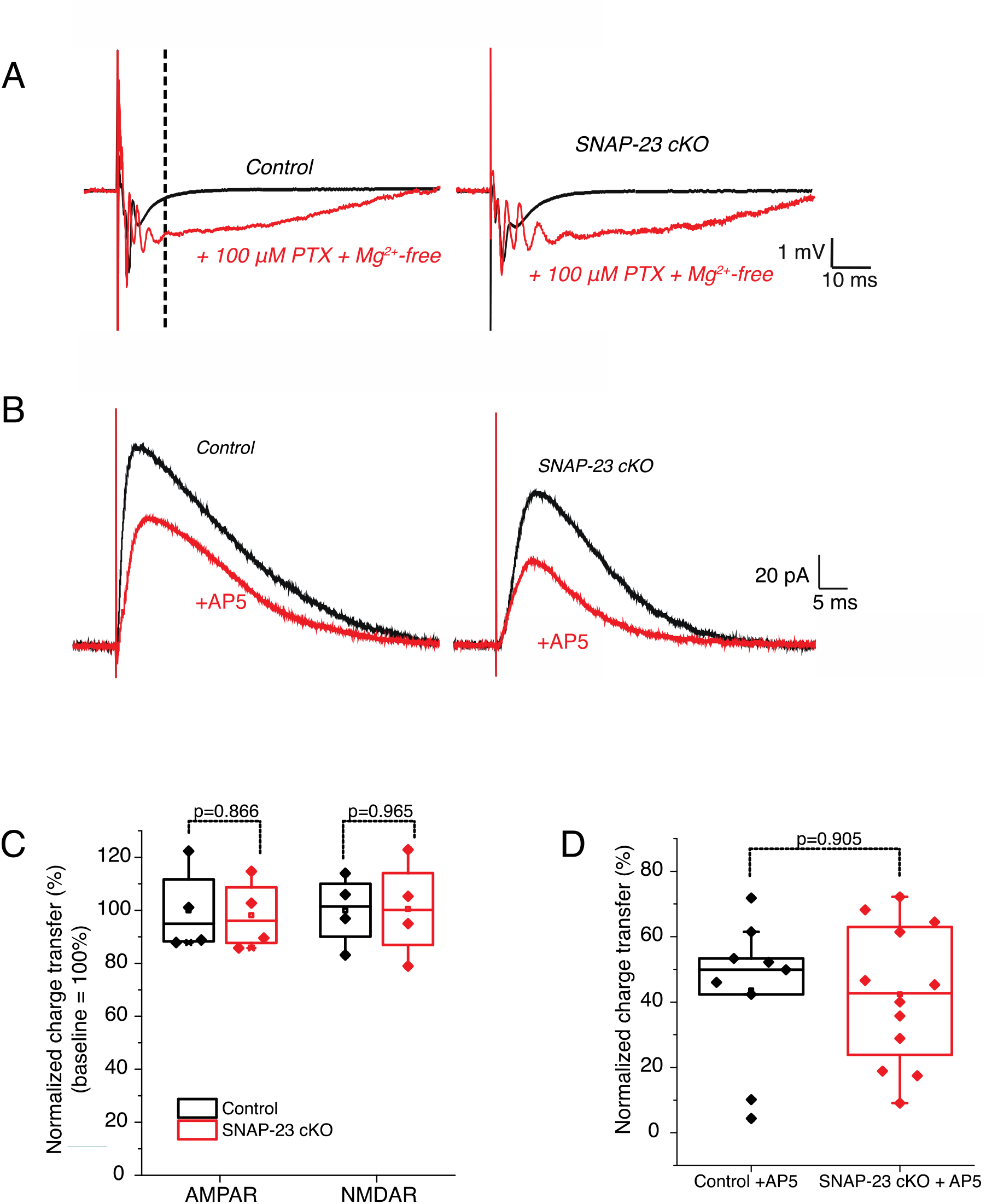
SNAP-23 is dispensable for AMPAR and NMDAR-mediated fEPSP and evoked NMDAR response. (A) Dendritic fEPSP recordings from the CA1 area of control and SNAP23 cKO mice in standard ACSF and in Mg2+-free ACSF plus 100 μM PTX (picrotoxin). Dashed line denotes from where AMPAR current calculation end and NMDAR current calculation begin. Stimulation artefact truncated for illustration purpose. **(B**) Representative traces of voltage clamp experiments where CA1 pyramidal neurons were voltage clamped at +40mV to observe both AMPAR and NMDAR-mediated responses. Recordings were done in 100 μM PTX containing ACSF. AP5 was bath applied for 5 minutes to block the NMDAR component so only the AMPAR-mediated component remains (red). (C) Normalized percent AMPAR and NMDAR-mediated charge transfer in control and SNAP23 cKO mice. The recordings obtained using Mg2+-free ACSF plus 100 μM PTX were analyzed for this purpose. The area under the curve during this AMPAR activity was calculated as AMPAR-mediated charge transfer while remaining response was calculated as NMDAR-mediated charge transfer. The charge transfer of both AMPAR and NMDAR was then normalized to the control group. Error bars indicate SEM (slice n= 4 for both groups, animal n = 3 for both groups). No significant group differences observed in AMPAR (p=0.866) or NMDAR (p=0.965) responses. Stimulation artefacts truncated for illustration purpose. (D) Normalized percent NMDAR charge transfer in control and SNAP-23 cKO mice. The area under the curve was calculated, the difference after AP5 application being the NMDAR component. The charge transfer was then normalized to the control group. No difference was observed in the SNAP-23 cKO mice compared to the control mice (p=0.905). Error bars indicate SEM.

To further investigate the effects of SNAP-23 deletion on the AMPA and NMDA currents, we performed whole-cell patch clamp recordings in CA1 neurons of the hippocampal slices (Figure 3B). CA1 neurons were voltage clamped at +40 mV to observe both AMPAR and NMDAR-mediated responses. After a stable baseline was obtained, AP5 was bath applied for 5 minutes, blocking the NMDAR component so only the AMPAR-mediated component remained. The area under the curve was calculated, the difference after AP5 application being the NMDAR component. AP5 application induced a 56.5% decreased in control slices, similarly, a 57.6 % decrease was observed in SNAP-23 cKO slices. Therefore, in agreement with our extracellular recording data, we found that NMDA currents were not changed in SNAP-23 cKO pyramidal neurons relative to controls (Figure 3D). Together, these data suggest that SNAP-23 does not play a role in regulation of basal AMPA or NMDA currents.

### SNAP-23 deficient CA1 neurons exhibit normal excitability

We then investigated whether the intrinsic excitability of the CA1 neurons is affected by SNAP-23 deletion. For this, current clamp was performed where constant square pulses (1 second) of increasing amounts of currents (from -300 pA to +500 pA in 25 pA increments) were injected and changes in voltage responses were monitored to gage the intrinsic excitability of CA1 neurons (Figure 4A). Single spike parameters of CA1 neurons were also measured using current clamp where shorter constant square pulses (100 ms) of increasing amounts of currents (from - 100 to +225pA in 25 pA increments) were injected and corresponding changes in voltage responses were monitored (Figure 4E). We found that the CA1 neurons of the SNAP-23 cKO mice exhibited similar basal intracellular properties and spiking properties as the control mice. The resting membrane potential, inter-spike intervals, spike half width and peak amplitudes were also similar between the two genotypes (Figures 4B–4D, 4F–4G).

**Figure 4.**
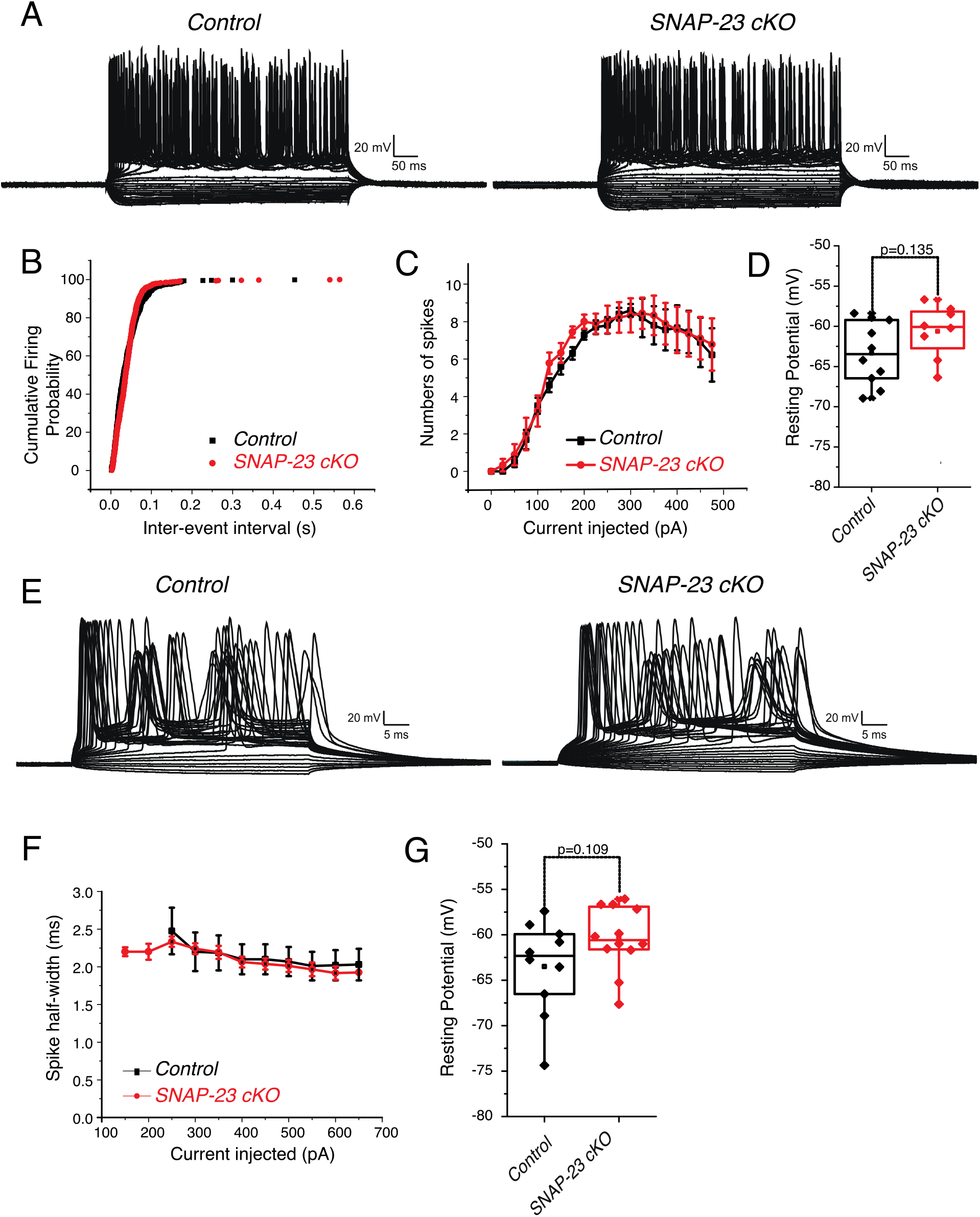
Intrinsic excitability remains the same in CA1 pyramidal neurons of SNAP-23 cKO mice. (A) Representative traces of current clamp experiments where currents from -300 pA to +500 pA were injected in 25 pA steps and changes in voltages from CA1 neurons of control and SNAP23 cKO mice were monitored. Intrinsic properties including cumulative firing probability (B), numbers of spikes induced by incremental current intensities (C) and resting membrane potentials (D). No difference was observed in membrane potential (p=0.135). (E) Representative traces of current clamp experiments where currents from -100 pA to +225 pA were injected in 25 pA steps and changes in voltage from CA1 neurons of control and SNAP23 cKO mice were monitored. Intrinsic properties including spike half width (F) and resting membrane potential (G). No difference was observed in resting membrane potential (p=0.109). Intrinsic properties were measured from neurons that had formed series resistance of < 20 MΩ for more than 10 min and resting membrane potentials less than −55 mV. Error bars indicate SEM.

We then monitored the excitatory postsynaptic currents (EPSCs) and examined the interevent intervals as well as amplitudes from individual CA1 neurons in the acute hippocampal slices. Using a holding potential of −70mV, we measured the frequencies and amplitudes of spontaneous EPSCs in CA1 neurons (Figure 5). Similar to evoked fEPSPs, we found that the average frequency and amplitude of the spontaneous EPSCs in the CA1 neurons remained unchanged in the SNAP-23 cKO mice (Figure 5D). Collectively, these data indicated that the postsynaptic responses observed in the SNAP-23 cKO CA1 neurons are unchanged in their basal properties and excitability and SNAP-23 removal does not compromise postsynaptic receptor currents.

**Figure 5.**
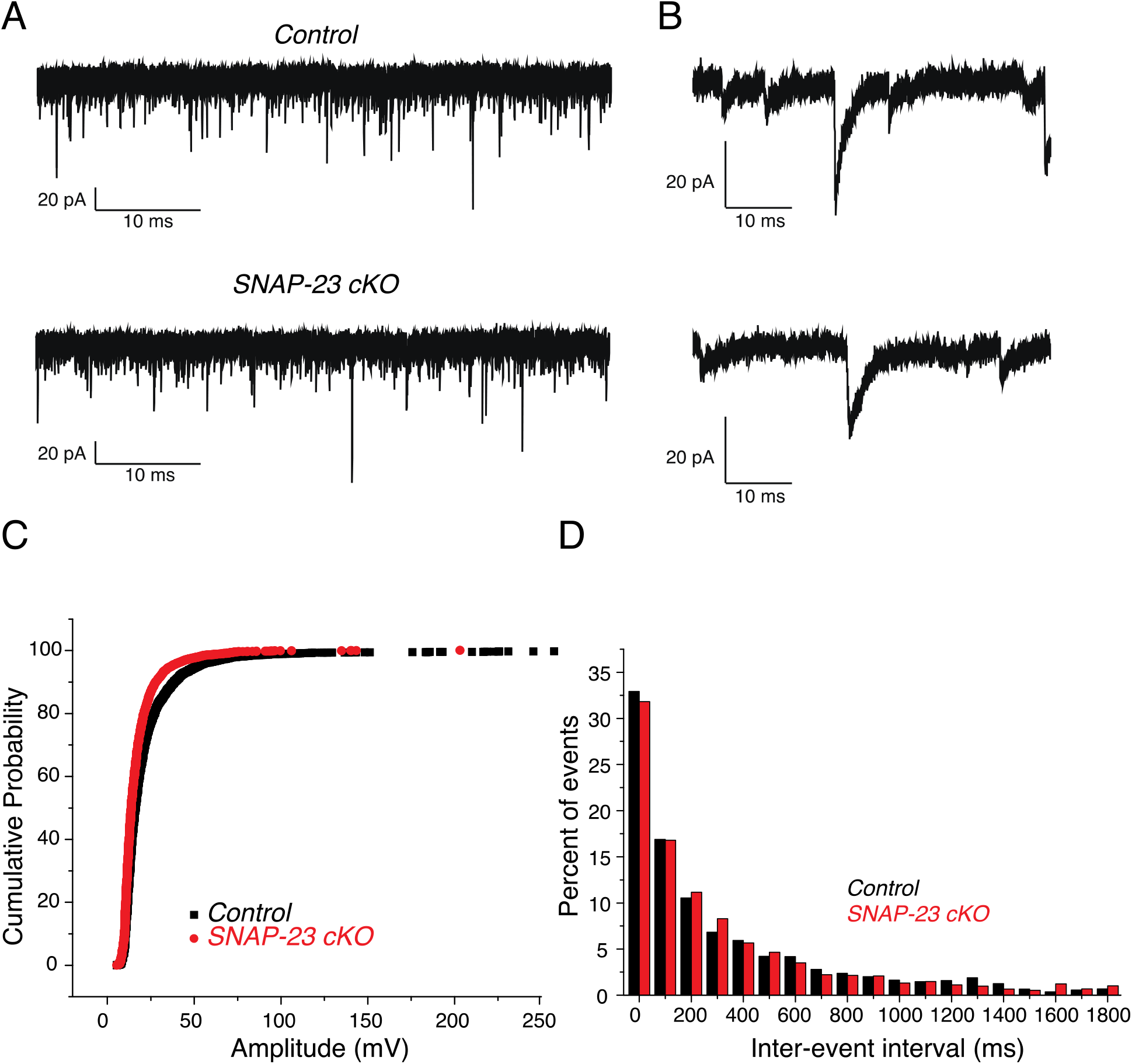
Amplitude and interevent interval of spontaneous EPSC of CA1 pyramidal neurons are not significantly altered in SNAP-23 cKO mice. (A) Representative traces of spontaneous EPSC recorded at −70 mV from CA1 pyramidal neurons of control and SNAP-23 cKO mice. (B) Expanded events as denoted in A. Cumulative amplitude (C) and inter-event interval (D) plot of spontaneous EPSCs of control and SNAP23 cKO. Data were collected from 9 control and 8 cKO CA1 pyramidal neurons. Spontaneous EPSCs were measured from neurons that had formed series resistance of < 20 MΩ for more than 10 min. Spontaneous EPSC that had amplitude less than noise level (8 pA) was omitted from analysis.

### SNAP-23 is essential for long-term potentiation

Having established that the SNAP-23 deletion does not affect basal hippocampal transmission, we next investigated the role of SNAP-23 in synaptic plasticity. Specifically, we tested whether SNAP-23 deletion affects long term potentiation (LTP), a major form of synaptic plasticity and a local circuitry correlate of learning and memory. For this purpose, we prepared acute hippocampal slices and measured the apical dendritic fEPSPs from CA1 before and after inducing LTP. We chose to use the theta-burst stimulation protocol as this is a more physiologically relevant LTP protocol compared to high-frequency stimulation. We gave 3 seconds of continuous theta-burst stimulation: 15 bursts of four pulses at 100 Hz, with an interburst interval of 200 ms in Schaffer collateral axons. We compared the magnitudes of post-theta stimulation potentiation (PTP) as well as the later LTP maintenance phase between control and SNAP-23 cKO mice. For each slice, the stimulation intensity was chosen such that it elicited 30% of the maximum response.

We examined CA1 LTP induction in 9 slices of 5 cKO mice and 8 slices from 4 control mice. Following the theta-burst stimulation delivery, PTP amplitudes and slopes being more than 150% of the baseline were observed for both control and the SNAP-23 cKO groups (Figures 6A-6C, supplemental Figure 2). There were no group differences in PTP measures [independent t-test, t_(15)_ = 0.62, p = 0.547; Figure 6C], In the control group, the PTP response transitioned to an evident LTP phase which stabilized at more than ~150% of baseline level and lasted as long as 60 minutes after the theta-burst stimulation (Figures 6A–6C). In contrast, the SNAP-23 cKO group showed a weak LTP phase which maintained at a level of less than 125% of baseline responses and was significantly diminished from the control [t_(15)_ = 3.83, p = 0.002]. These observations, together with the above assessments of CA1 evoked fEPSPs (Figure 2), indicate that SNAP-23 is selectively critical for synaptic plasticity without affecting basal transmission.

**Figure 6.**
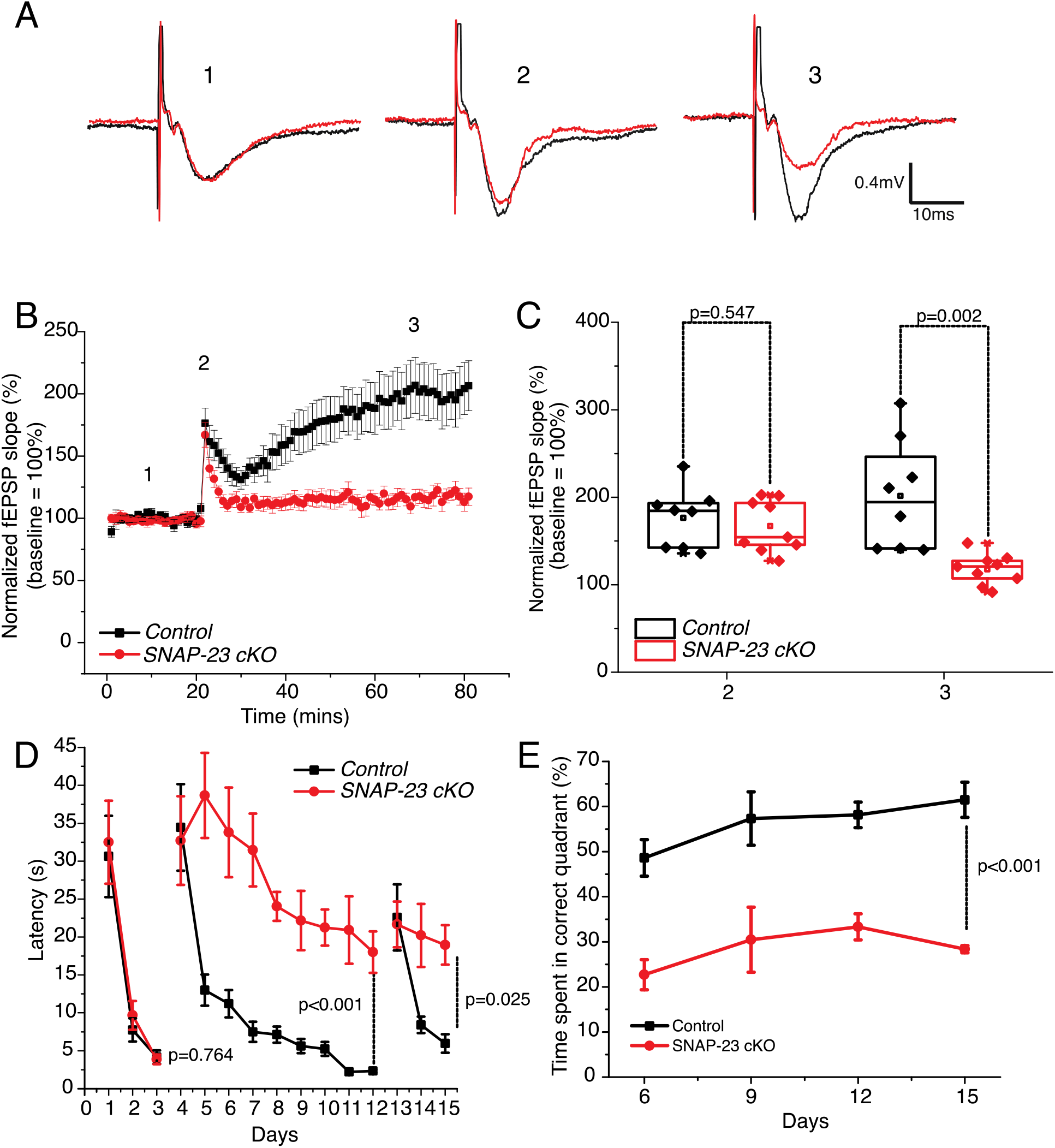
CA1-specific SNAP-23 cKO results in defective Schaffer collateral-CA1 LTP and significantly impairs Morris water maze performance. (A) Representative recordings of fEPSP of control and SNAP-23 cKO CA1 at baseline (1), immediately after LTP induction (2), and 50 min after LTP induction (3). Each illustrated trace was averaged from four consecutively evoked responses. (B) average field EPSP slope normalized against baseline. Data were collected from 9 slices of KO mice and 8 slices of control mice. (C) Normalized field EPSP slope control and SNAP-23 cKO at times 2 (immediately after LTP induction) and 3 (50 mins after LTP induction). Error bars indicate SEM (p=0.546 for timepoint 2, p=0.002 for timepoint 3). (D) learning curves for mice to find the visible platform (days 1-3), hidden platform (days 4-12) and hidden platform during reversal training (days 13-15). Animals were tried 4 times a day, with randomized entry points to the pool. Latency to find the hidden platform was significantly increased in the SNAP-23 cKO mice relative to the controls, however SNAP-23 cKO mice still demonstrate some degree of learning as determined by a mixed ANOVA test. (E) During Probe Test, platform was removed from the pool and % of time spent in quadrant where platform was previously located was calculated from total recording time of 60 sec. Error bars indicate SEM, two-sample t-test.

### Tissue-specific SNAP-23 KO mice exhibit impaired spatial memory

We next investigate the behavioral consequences of LTP deficits seen in the SNAP-23 cKO mice by performing Morris water maze to assess the spatial learning and memory (31, 32). A 15-day protocol was performed where each day was composed of four separate trials. During the four trials, the mice were placed into the pool from a randomized cardinal position (Figure 6D-E). The first three days, a flag was present on the platform as a visible cue to allow mice to learn the cued task. Day 4 onwards, the flag was removed, the position of the hidden platform was changed, and the mice were asked to learn the position of the hidden platform. The SNAP-23 cKO group (n = 6) learned the visible cued swimming task as latency to find the platform were similarly reduced to control group (n = 6) by day 3. There was no statistical difference between the two groups [Mixed ANOVA: F_(1,10)_ = 0.095, p = 0.764], indicating that the SNAP-23 cKO did not impair the ability of to visualize surroundings and swim (Figures 6D and 6E). From day 4, the mice were tasked to learn the position of a hidden platform. The control animals had a steep drop in the latency to find the platform between days 4 and 5, quickly learning the new location of the platform demonstrating effective spatial learning. On the other hand, the SNAP-23 cKO had impaired spatial learning and memory as mice took significantly more time to find the hidden platform. Mixed ANOVA showed that there is a statistically significant difference between the two groups [F_(1,10)_ = 101.9, p < 0.001]. SNAP-23 cKO did exhibit some learning capacity as a decrease was seen in the latency by day 12 of the trial [effects of days: F_(8,10)_ = 14.752, p < 0.001], but this learning was at a significantly slower rate to that of the littermate controls [interactions between groups and days: F_(1,8)_ = 3.992, p < 0.001]. SNAP-23 cKO mice also spent less time in the correct quadrant during probe tests (Figure 6E) further demonstrating the deficit in spatial learning from the SNAP-23 deficiency [Mixed ANOVA: F_(1,10)_ = 55.420, p < 0.001]. On day 13, a reversal training was instated, and the location of the hidden platform was again changed. On day 13, both the control and SNAP-23 cKO groups needed more time to find the platform. However, the control group was able to re-learn the location more quickly than the SNAP-23 cKO group [F_(1,10)_ = 6.924, p = 0.025]. Together, our results indicate that SNAP-23 in the CA1 neurons of the hippocampus is critical for spatial memory.

### The behavioral abnormality of SNAP-23 cKO mice is more selective than that of syntaxin-4 cKO mice

We previously found that in syntaxin-4 cKO mice, basal transmission as well as LTP were both strongly decreased (25), which supports previous findings that syntaxin-4 directs synaptic plasticity (33). Like the SNAP-23 cKO, the syntaxin-4 cKO mice showed deficits in their performances of the Morris water maze (25). Here, we investigated other sets of behavior tasks using the SNAP-23 and the syntaxin-4 cKO mice to uncover any correlations between synaptic and behavioral abnormalities. We first performed marble burying test and nest shredding test (Figure 7A-B). Marble burying and nestlet shredding are tasks often used to gain insight into repetitive behaviors, moreover, nestlet shredding is a task that involves the hippocampus (34). We found that the marble burying was unchanged in the syntaxin-4 cKO mice (n = 6) compared to the littermate controls (n = 6) [independent t-test, t(10) = −0.31, p = 0.765]. However, the syntaxin-4 cKO mice showed differences in the nestlet shredding task, completing the test with less amounts of shredded nestlet [height t_(10)_ = 5.59, p = 0.001; volume t_(10)_ = 6.50, p=0.001]. In contrast, there was no change to the level of repetitive behavior of the SNAP-23 cKO mice in both tests (Figure 7A).

**Figure 7.**
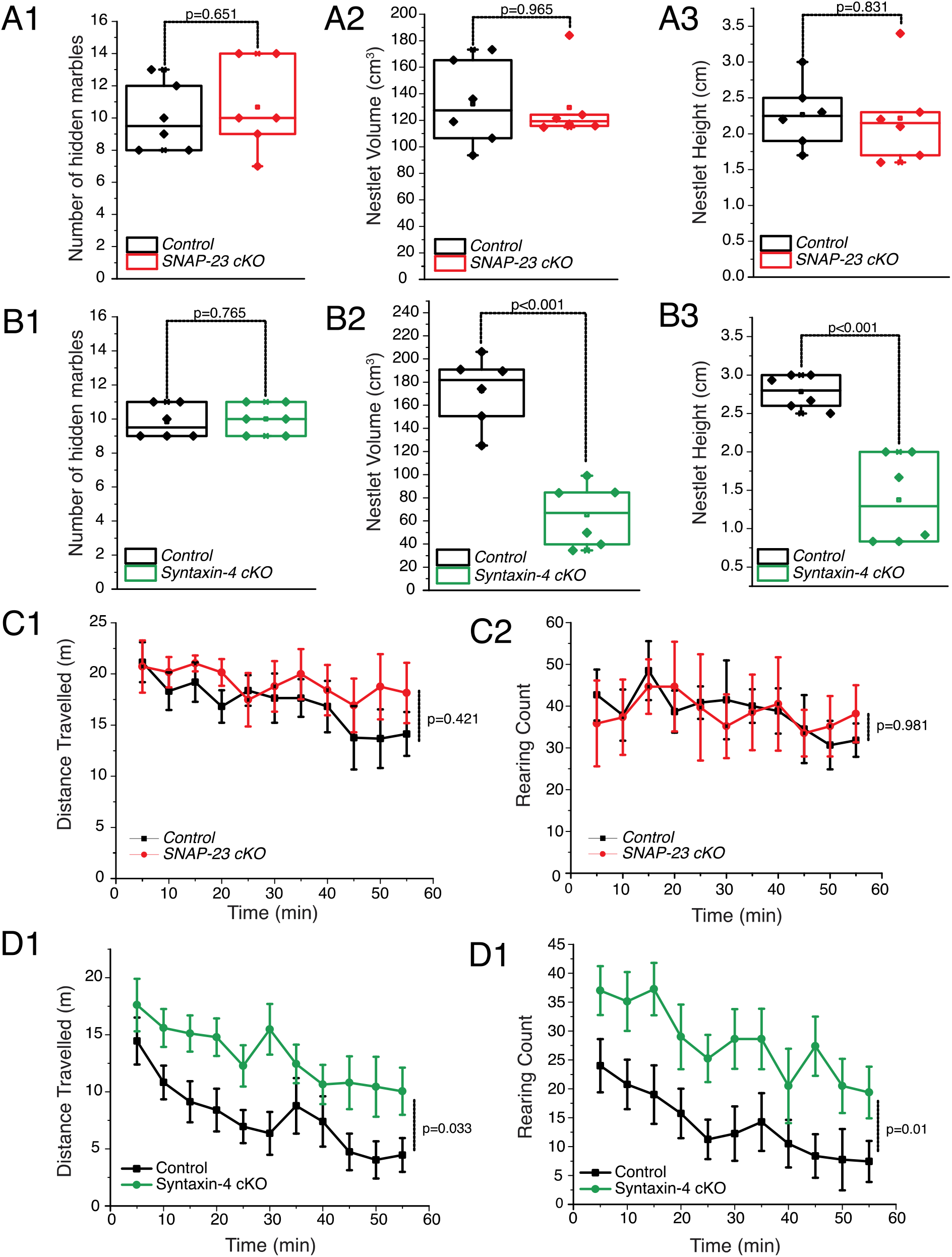
SNAP-23 cKO impairment is memory specific leaving other behaviors intact while Syntaxin-4 cKO impairs other behaviors such as nestlet shredding. Quantifications of marbles buried and nestlet shredded by SNAP-23 cKO (A) and syntaxin-4 cKO mice (B); both mice buried a comparable number of marbles when compared to control (p = 0.651 SNAP-23 cKO; p = 0.765 for syntaxin-4 cKO). However, syntaxin-4 cKO mice shredded less amount of nestlet compared to control (p = 0.001 for nestlet height; p = 0.001 for nestlet volume) while SNAP-23 shredded amounts similar to that of control (p = 0.830 for nestlet height; p = 0.96 for nestlet volume; two sample t-test). Open field test of SNAP-23 (C) and syntaxin-4 (D) mice, distance travelled and rearing count were assessed. SNAP-23 cKO mice show no changes to both distance travelled or rearing count when compared to littermate controls, while syntaxin-4 cKO mice show an increase to spontaneous locomotion and rearing count when compared to littermate controls. Error bars denote SEM, mixed ANOVA test.

We also performed an open-field test to assess changes to the level of anxiety in these animals. Here, we evaluated spontaneous locomotor activity and the rearing pattern of each genotype under a novel environment (Figure 7C-D). We found that syntaxin-4 cKO strikingly increased in spontaneous locomotion as well as rearing behavior compared with littermate control mice. However, we did not find difference in these between SNAP-23 cKO and the littermate control group. These results together suggest that changes in basal neurotransmission seems to affect repetitive and spontaneous locomotor activity. However, LTP is more specifically required for learning and memory wherein SNAP-23 plays a crucial role.

## Discussion

By generating a pyramidal neuron-specific knockout of SNAP-23, we were able to demonstrate the role of neuronal SNAP-23 in basal synaptic transmission and synaptic plasticity as well as its behavioral consequence. Deletion of SNAP-23 did not show major abnormalities in the morphology of the hippocampus (Figure 1), basal synaptic transmission (Figure 2) including AMPAR and NMDAR mediated postsynaptic currents (Figure 3, 5), intrinsic excitability of CA1 pyramidal neurons (Figure 4) or repetitive and spontaneous locomotive behavior (Figure 7). However, SNAP-23 cKO impaired LTP and spatial memory (Figure 6). Therefore, we conclude that neuronal SNAP-23 is selectively crucial for synaptic plasticity and spatial memory.

Studying the mechanisms of receptor trafficking is difficult due to the fact that neurons express multiple isoforms including the neuronal SNAP 25, but also SNAP-23 (16, 21, 35), SNAP-29 (36), and SNAP 47 (19, 37, 38). The expression of these isoforms may suggest for distinct roles, however, it also introduces confound that in the absence of one protein, one or more of the other isoforms may compensate for the loss. Indeed, it has been shown that SNAP-47 can substitute for SNAP-25 in cultured neurons, albeit with less efficacy (38). As such, previous publications have reported highly contradicting results. Such lack of consensus may also be due to differences in methodologies taken to study these proteins, as a lethality is observed in both SNAP-25 (39) and SNAP-23 (26) global knockout mice. In fact, even a broader and milder nervous system conditional knockout of SNAP-23 using Nestin-Cre which is expressed in neuronal and glial cell precursors resulted in severe phenotypes with animals dying within 3 weeks of birth (40). These Nestin-Cre induced SNAP-23 knockout mice lack hippocampi, further emphasizing the importance of SNAP-23 in brain development. Therefore, other studies on SNAP-23 as well as other SNAP isoforms have largely relied on transient shRNA knockdown (21, 22, 41), and heterozygote (21) approaches. SNAP-47 has been suggested to be involved in AMPAR trafficking (22), while SNAP-23 and SNAP-25 have been suggested to be involved in NMDAR trafficking (21, 42). One group found that SNAP-23, and not SNAP-25, is responsible for regulating NMDAR expression (21). However, also using shRNA knockdown, another group reported different results and found that SNAP-25, and not SNAP-23, was important for AMPAR trafficking during LTP (22). This study also found that SNAP-25, not SNAP-23, regulates surface NMDA receptor levels and reduction of SNAP-23 fails to impair NMDAR-dependent chemical LTP (22). However, a later review brings up the key point that SNAP-23 may play a more developmental role and/or residual SNAP-23 from the incomplete knockdown may be sufficient in supporting LTP (36). Finally, another group described SNAP-25, but not SNAP-23, being responsible for enhancing NMDA currents by NMDAR delivery (41). While shRNA knockdowns were indeed able to show changes in glutamate receptor trafficking (21), or changes to LTP in the case of SNAP-47 (22), studying the behavioral consequences of these findings is a challenge and difficult due to lack of viable animal models.

With pyramidal neuronal-specific SNAP-23 cKO mice, our data provide evidence that SNAP-23 does not appear to be required in supporting baseline AMPAR and NMDAR responses in CA1 pyramidal neurons, but selectively mediates LTP. Previous studies suggested that SNAP-47 plays this role (22). The discrepancies between our and previously reported results (21, 22) can be due to various reasons, such as chronic SNAP-23 deletion causing compensation by other isoforms, or transient knockdowns being not sufficiently removing the protein of interest. Non-specific targeting of shRNA can also contribute to such contradicting results. However, as we observe in our model, despite the chronic removal of SNAP-23 not affecting AMPAR and NMDAR currents, it was able to impair LTP and spatial memory and we were able to show, for the first time, behavioral consequences arising from removal of SNAP-23. As these impairments to LTP and spatial memory were observed in adult mice, it also challenges the notion that SNAP-23 does not play a role in supporting NMDAR currents in adulthood (36).

Unlike what we previously observed in syntaxin-4 cKO mice, in which both basal neurotransmission and LTP were diminished, SNAP-23 cKO mice are only impaired in LTP while basal neurotransmission remains intact. While syntaxin-4 cKO mice exhibit abnormalities in various behaviors, deficits of SNAP-23 cKO is only limited to spatial memory. Syntaxin-4 cKO mice showed abnormalities in the nestlet shredding test, which is contrast to SNAP-23 mice where the performance of nestlet shredding was comparable to that of the control. This suggests that the LTP deficits are spatial learning and memory specific as demonstrated by poor Morris water maze performances, while deficits to the basal neurotransmission may affect other behaviors such as the hippocampus-dependent nestlet shredding task. The observed SNAP-23 cKO phenotypes are also different from that of syntaxin-3 cKO, as syntaxin-3 cKO led to no changes in LTP or basal neurotransmission as well as no changes in learning and memory (29). This suggests that the regulation of glutamate receptor trafficking in postsynaptic membranes are highly heterogenous, and requirements of specific SNARE proteins may depend on the neuronal activity state.

Previously, we proposed that syntaxin-4 may play a direct role in ionotropic glutamate receptor trafficking, and is responsible for maintaining basal neurotransmission, synaptic plasticity and hippocampus-dependent learning; syntaxin-4 cKO leads to decreased basal ionotropic glutamate receptor levels and AMPAR insertion during LTP thereby impairing basal neurotransmission and LTP. We also speculate that syntaxin-4 exerts an indirect effect – perhaps the decrease in basal NMDAR current observed in syntaxin-4 cKO results in decreased calcium influx, not allowing for sufficient activation of the signaling cascade that induces AMPAR insertion during LTP. In this model, activity-dependent AMPAR trafficking would depend on t-SNAREs syntaxin-4, and as well as SNAP-23 which we elucidated in this study albeit through different mechanisms of action (Figure 8). Under this model, SNAP-23 cKO does not alter basal neurotransmission, and only alters activity-dependent AMPAR delivery downstream of the calcium signaling cascade resulting in deficits to synaptic plasticity.

**Figure 8.**
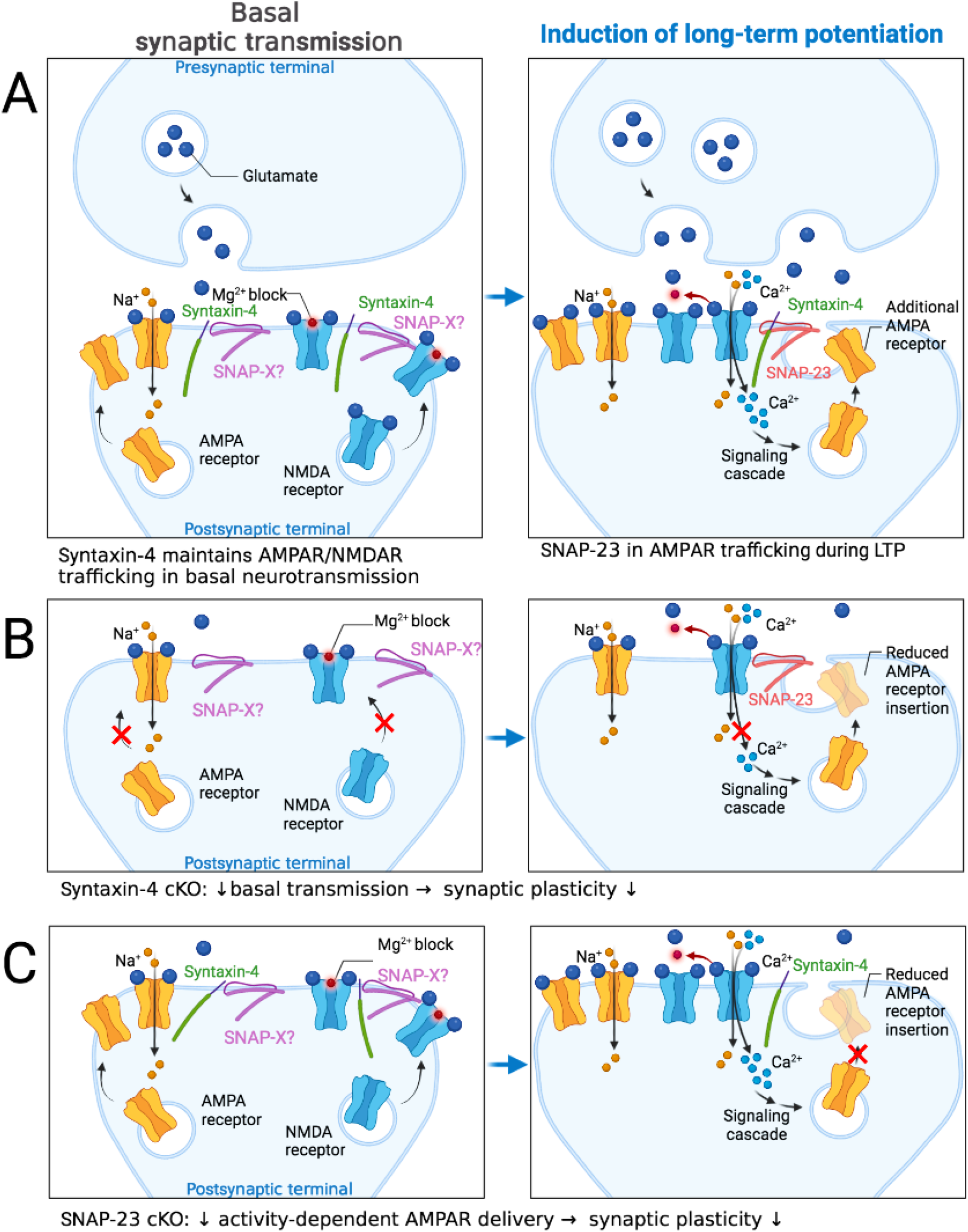
Model of SNAREs involved in basal transmission and LTP. (A) Left panels indicate the presynaptic and postsynaptic terminal under basal synaptic transmission conditions. t-SNAREs syntaxin-4 and an unidentified SNAP isoform (SNAP-X) maintain AMPAR and NMDAR levels under basal synaptic transmission conditions. Right panel indicates synaptic changes following induction of long-term potentiation (LTP); following removal of the magnesium block by theta burst stimulation, calcium influx activates signaling cascades to trigger activity-dependent AMPAR insertion. Activity dependent exocytosis requires syntaxin-4 complexing with SNAP-23 to deliver AMPARs during LTP. (B) Syntaxin-4 cKO decreases basal AMPAR and NMDAR numbers and decreases AMPAR insertion during LTP. Also possible is an indirect effect of syntaxin-4 cKO decreases basal NMDAR levels, allowing for less calcium influx during LTP, and reducing AMPAR insertion. (C) SNAP-23 cKO does not alter basal neurotransmission, and acts downstream of calcium signaling cascade, reducing AMPA receptor insertion during LTP. Adapted from “Long-Term Potentiation”, by BioRender.com (2022). Retrieved from https://app.biorender.com/biorender-templates.

In summary, for the first time, our SNAP-23 cKO mice allowed us to test the role of SNAP-23 *in vivo* and *in vitro*. Conditional knockout mice survived to adulthood and allowed spatial learning and memory tests to be performed, opening up new potentials for a model of Alzheimer’s disease (43). Our results using the Morris water maze task indicate that the reduction of LTP maintenance is correlated as deficits in spatial learning and memory (Figures 6). However, AMPAR and NMDAR currents were not reduced, and NMDA evoked responses were not reduced, therefore suggesting a dispensable role of SNAP-23 in glutamate receptor trafficking during basal neurotransmission.

## Experimental Procedures

### Animals

We used animals with an age between 2.5 - 6 months exclusively for all experiments without preferences on sex. The mice were housed in a vivarium that was maintained between 22 - 23°C with a 12-hr light on/off cycle. Food and water were accessible *ad libitum*. All experiments detailed here were reviewed and approved by the animal care committee of the University Health Network in accordance with the Canadian Guidelines for Animal Care.

### Generation of tissue-specific SNAP-23 cKO mice

SNAP-23 flox/flox mice were previously created by OzGene by introducing a loxP site flanking exons 3 and 4 (24). CaMKIIα-Cre mice were purchased from Jackson Laboratory (005359). SNAP-23 flox/flox mice were crossed with CaMKIIα-Cre to generate tissue-specific SNAP-23 conditional KO mice. For each animal, genomic DNA was extracted from the tail for genotyping by PCR. SNAP-23 conditional KO mice were viable, fertile and developed to adulthood without obvious behavioral abnormalities or welfare concerns. SNAP-23 flox/flox mice without CaMKIIα-Cre were used as control mice for all experiments. One possible caveat of our model is that although CaMKIIα-Cre is generally accepted as a forebrain specific Cre recombinase line, it has also been previously shown that CaMKIIα-Cre can be minimally expressed in the male testis (44). This introduces a small possibility of a global ablation to SNAP-23. To mitigate this confound, breeding pairs were set up so the CaMKIIα-Cre positive parent was the female to prevent transmission of Cre recombinase through male germ cells.

### PCR for genotyping

Mouse tails were collected, and genomic DNA was obtained by alkaline lysing methods where the samples were lysed with 50 mM NaOH while incubating at 96 °C for 1 hour with vigorous shaking. The supernatant was directly used in PCR for genotyping. PCR was performed to distinguish wild-type vs flox mice and presence of Cre recombinase was also tested. For 5’ PCR, a forward primer 5’-GGGGGTGAGTTGAAGTCATTGAAG-3’ and a reverse primer 5’-AGCTTAAACGGGATGAACTCAGGC--3’. Primers are located on the end of the end of exon 4 and near the start of intron 4, surrounding the closing loxP site. For Cre recombinase PCR, the forward primer of 5’-CCCCAAGCTCGTCAGTCAA and the reverse primer of 5’-ACAGAAGCATTTTCCAGGTATGCT-3’ were used.

### Preparation of hippocampal sections for two photon imaging

Mice were anesthetized with an intraperitoneal injection of sodium pentobarbital (75 mg/kg, Somnotol, WTC Pharmaceuticals) and transcardially perfused with 10 mL of PBS followed by 10 mL of 4% PFA. The brain was removed and postfixed overnight at 4°C in 4% PFA. The brain was then washed in PBS and dehydrated in 30% sucrose overnight. Brains were mounted in 50% OCT compound and 50% sucrose and 40 μm coronal sections were prepared using a cryostat.

For immunohistology, slices were washed in PBS with 0.1% triton X-100. Rhodamine-phalloidin was incubated for 90 mins. After washing, slices were then blocked for one hour in PBS with 0.3% triton X-100 and 10% goat serum. Primary antibody was incubated overnight in blocking solution. Primary SNAP-23 antibody is from Novus Biologicals (JA73-15, Cat. NBP2-67157) used at 1:125 dilution. After washing, secondary antibody (A-11008) Alexa Fluor 488 Goat anti-Rabbit IgG) was incubated for 2 hours in blocking solution at 1:1000 dilution. Slices were then washed in PBS with 0.1% triton X-100 and mounted in Mowiol.

### Two-photon fluorescence imaging of CA1 hippocampus in hippocampal sections following anti-SNAP23 & phalloidin co-staining

Whole-brain coronal sections were prepared as described above. Fluorescence imaging of the CA1 hippocampus was conducted with a two-photon microscope (Custom made FV1000 MPE; Olympus, Tokyo, Japan, 60x objective lens, NA 1.0 equipped with Spectra-Physics InSight DeepSee). An 860 nm laser was used for excitation. Green (Alexa Fluor 488 dye) and red (phalloidin) fluorescence were imaged at 495-540 nm and 575-630 nm emission, respectively. The fluorescence image was combined across a z-stack composed of 6 slices, taken at 0.5 μm intervals for each slice.

### Cresyl violet staining of hippocampal sections

Mice were anesthetized and prepared as described above, however transcardial perfusion was done with a 10% neutral buffered formalin solution (Sigma, Oakville, Canada). The brain was removed, postfixed, mounted, and a series of coronal cryosections were obtained at 40 μm thickness and stained with cresyl violet. Slices were imaged on a OMAX A3580U camera on a dissecting microscope with the OMAX ToupView v3.7 software. Quantification of hippocampus area and density was done in ImageJ, briefly, the hippocampus was manually selected and area and mean gray intensity were measured as a function of area and intensity respectively.

### Preparation of hippocampal slices for electrophysiological recordings

Mice were anesthetized using an intra-peritoneal injection of sodium pentobarbital (75 mg/kg) and transcardiacally infused with ice-cold high sucrose dissection solution containing: 300 mM sucrose, 3.5 mM KCl, 2 mM NaH_2_PO_4_, 20 mM glucose, 0.5 mM CaCl_2_, 2 mM MgCl_2_ and 5 mM HEPES (pH adjusted to 7.4) before decapitation. The brain was quickly dissected and hemi-sectioned, and mid-line sides of two hemispheres of the brain were glued. Sagittal cortico-hippocampal slices of 400 μm thickness were obtained via a vibratome (VT1200, Leica Microsystems, Richmond Hill, Canada) in the presence of ice-cold high sucrose dissection solution and cortical tissues were removed leaving only the hippocampus. For each animal, an average of 10 - 12 hippocampal slices were sectioned. After sectioning, the slices were stabilized in artificial cerebrospinal fluid (ACSF) containing: 125 mM NaCl, 25 mM NaHCO_3_, 3.5 mM KCl, 1 mM NaH_2_PO_4_, 1 mM MgSO_4_, 2 mM CaCl_2_ with continuous gassing with 95 % O_2_ and 5 % CO_2_ at room temperature for at least 1 hour before the recording.

### Electrophysiological recordings

The slice was placed in a submerged chamber and perfused with standard, oxygenated ACSF at a high flow rate of 15 mL/min. Both the top and bottom surfaces of the slice were exposed to ACSF. All recordings were done at room temperature. For all recordings, recording electrodes and stimulating electrodes were positioned under a dissecting microscope.

Recording electrodes were made with thin-wall glass tubes (World Precision Instruments, Sarasota, Florida). Extracellular electrodes were filled with a solution that contained 150 mM NaCl and 2 mM HEPES (pH 7.4; resistance of 1 - 2 MΩ). Electrodes for whole-cell current-clamp recordings were filled with solution that contained 140 mM potassium gluconate, 1 mM HEPES, 1 mM EGTA, 0.1 mM CaCl2, 0.5 mM NaGTP, 5 mM MgATP, and 5 mM phosphocreatine (pH 7.26 and resistance of ~5 MΩ); electrodes for whole-cell voltage-clamp recordings were filled with solution that contained 120 mM cesium gluconate, 10 mM TEACl, 10 mM HEPES, 2 mM EGTA, 1 mM CaCl2, 1 mM MgCl2, 0.5 mM NaGTP, 5 mM MgATP, and 5 mM phosphocreatine (pH 7.18 and resistance ~5 MΩ).

Extracellular and intracellular signals were recorded via a dual channel amplifier (700B, Molecular Devices/Axon Instruments, Sunnyvale, California). Data acquisition, storage and analysis were performed using PClamp software (version 10, Molecular Devices). These signals were recorded in frequencies of 0 - 5 kHz and digitized at 50 kHz (Digidata 1322A, Molecular Devices). For assessing evoked synaptic field potentials, extracellular recordings were made from the apical dendritic area of the CA1. For assessing intrinsic properties and spontaneous EPSC, CA1 somas were recorded via whole-cell patch clamp. Intrinsic properties were assessed via current clamp. Intracellular injections of squared current pulses of -300 pA to 500 pA in 25 pA steps and duration of 500 ms or intracellular injections of square current pulses of100 pA to 225 pA in 25 pA steps and duration of 50 ms were used to assess firing properties and spike properties. A bipolar electrode, made of polyimide-insulated stainless-steel wires (outer diameter 0.1 mm; Plastics One, Ranoake, Virginia), was placed in the *stratum radiatum* CA2 region for Schaffer collateral axon stimulation. Constant current pulses were generated by a Grass stimulator (S88H, Natus Neurology Incorporated – Grass Products, Warwick, Rhode Island) and delivered through an isolation unit.

For spontaneous EPSC recordings, the neurons were held at −70 mV via voltage clamp. Spontaneous EPSCS were analyzed using MiniAnalysis (Synaptosoft). For these voltage clamp recordings, we used a cesium-gluconate-based pipette solution which showed pipette resistance of 5 MΩ in standard ACSF. Only cells that showed stable responses for 10 minutes after establishing a whole-cell configuration were used.

### LTP induction

Stimulation at which results in 30% of the maximal evoked response of fEPSP slopes was quickly determined by giving constant current pulses (10 - 150 μA, 1 ms). Then the stability of evoked fEPSPs was monitored for 30 min prior to LTP induction. Slices which exhibited deviations of more than 10 % in the baseline responses were rejected from further recordings.

LTP was induced by applying 3 sec of continuous theta burst stimulation: 15 bursts of four pulses at 100 Hz, with an interburst interval of 200 msec. After theta burst stimulation, responses were further recorded for 60 min.

### Pharmacological agents and blockade experiments

(2R)-amino-5-phosphonovaleric acid (AP5) was obtained from Sigma/Research Biochemicals Inc. (Mississauga, Canada), and picrotoxin was obtained from Tocris Bioscience (Bristol, United Kingdom). AP5 was initially dissolved in ddH2O to a stock concentration of 10 mM. The final concentration applied was 50 μM. Picrotoxin at 100 μM was added directly to the Mg^2+^-free ACSF and was dissolved with an extensive period of stirring. Briefly, baseline recordings were made under standard ACSF. Then, the bath solution was switched to Mg^2+^-free ACSF containing 100 μM picrotoxin while stimulating the Schaffer collateral axons every 30 sec to maximize glutamatergic activations and responses. Epileptiform responses with long lasting phase with multiple spikes started to appear after 15 min.

### Morris Water Maze Test

Mice received visible platform training for 3 days (4 trials per day) and hidden platform training for 12 days (4 trials per day). If the mice did not find the platform within 90 seconds, they were guided to the platform by the experimenter’s hand. For hidden platform training, the platform was submerged under 1.5 cm of water and the procedure is otherwise the same as visible platform training. The location of the platform was changed in hidden platform training compared to visible platform training. For reversal training, the location of the platform was changed to a different quadrant from the regular hidden platform training. On days 6, 9, 12 and

15, mice were given a 60 second probe trial where they were allowed to explore the maze without platform. The inter-trial interval was roughly 10 - 15 minutes. Learning was assessed by evaluating the time required to find the hidden platform in the training trials and memory was measured by examining time spent during the probe trials in the quadrant of the pool where the platform was previously located.

### Marble burying test

Marble burying task was performed with minor modifications to a previously published protocol (45). Briefly, cages were half filled with 5 cm of corn bedding, and 20 marbles were lined approximately 2 cm apart within the cage. Mice were each placed on the bedding in a corner of the cage, and the cage lid closed. Mice were left undisturbed and allowed to freely dig around the cage for an hour, food and water were withheld during the test. At the end of the session, any marble at least two-thirds covered with bedding were scored as buried.

### Nestlet shredding task

Nestlet shredding task was performed with minor modifications to a previously published protocol (45). Briefly, mice were placed in a cage with a single, preweighed nestlet, and the cage lid closed. Mice were left undisturbed and allowed to roam around the cage for an hour, food and water were withheld during the test. The mouse was returned to its home cage after test completion. Nestlet height (in cm) and nestlet volume (in cm^3^) were then measured.

### Open-field test

Open field ambulation test was performed with minor modifications to a previously published protocol (46). Briefly, mice were placed in a plexiglass cage for an hour. An automated movement detection system recorded the motor activities of the animal. Measured parameters were distance travelled and rearing count.

### Statistics

Statistics were performed using Origin Pro 2016 (OriginLab, Northampton, MA) and SPSS (IBM SPSS Statistics, Armonk, NY). All error bars represent SEM. For comparison of two groups, a two-sample t test was performed using Origin Pro. Mixed ANOVA testing was performed in SPSS. For all box plots, box-and-whisker plots represent the median (central line), 25th–75th percentile (bounds of the box), and outliers (whiskers)

## Author Contributions

M.H. and N-R.B. designed, performed and analyzed experiments and wrote the manuscript. K.M. contributed to electrophysiological recordings and behavioral tasks. J.R. and K.O. contributed to two-photon microscopy experiments. N-R.B., K.M., J.B.X, and D.F. assisted on generation of SNAP-23 cKO mice and maintenance of mouse colony. H.G. and J.P. advised on genotyping of mice. H.H. and C.H.C. assisted with histological experiments. S.E., H.S.S. and Z.P.F assisted with imaging experiments. L.Z. assisted on electrophysiological recordings on slices. S.E., H.S.S., Z.P.F., H.G., J.P., P.P.M., K.O., L.Z. and S.S. helped with editing the manuscript. S.S. supervised the project.

## Acknowledgements

This research was supported by the *Natural Sciences and Engineering Research Council* of Canada (RGPIN-2015-06438) and the Canadian Institute of Health Research (MOP-130573). J.R. and K.O. were supported by the Canadian Institute of Health Research (PJT 156103, PJT 165917) and the *Natural Sciences and Engineering Research Council* of Canada (RGPIN-2017-06444). M.H. and C.H.C. are supported by the Vision Science Research Program scholarship, and the Ontario Graduate Scholarship. The Alexander Graham Bell Canada Graduate Scholarships-Doctoral (CGS-D2) from Natural Sciences and Engineering Research Council of Canada was issued to N-R.B. 2017 Spring Postdoctoral & Clinical Research Fellowship was issued to N-R.B.

