## Supplemental Figures for "Neuronal SNAP-23 scales hippocampal synaptic plasticity and memory"

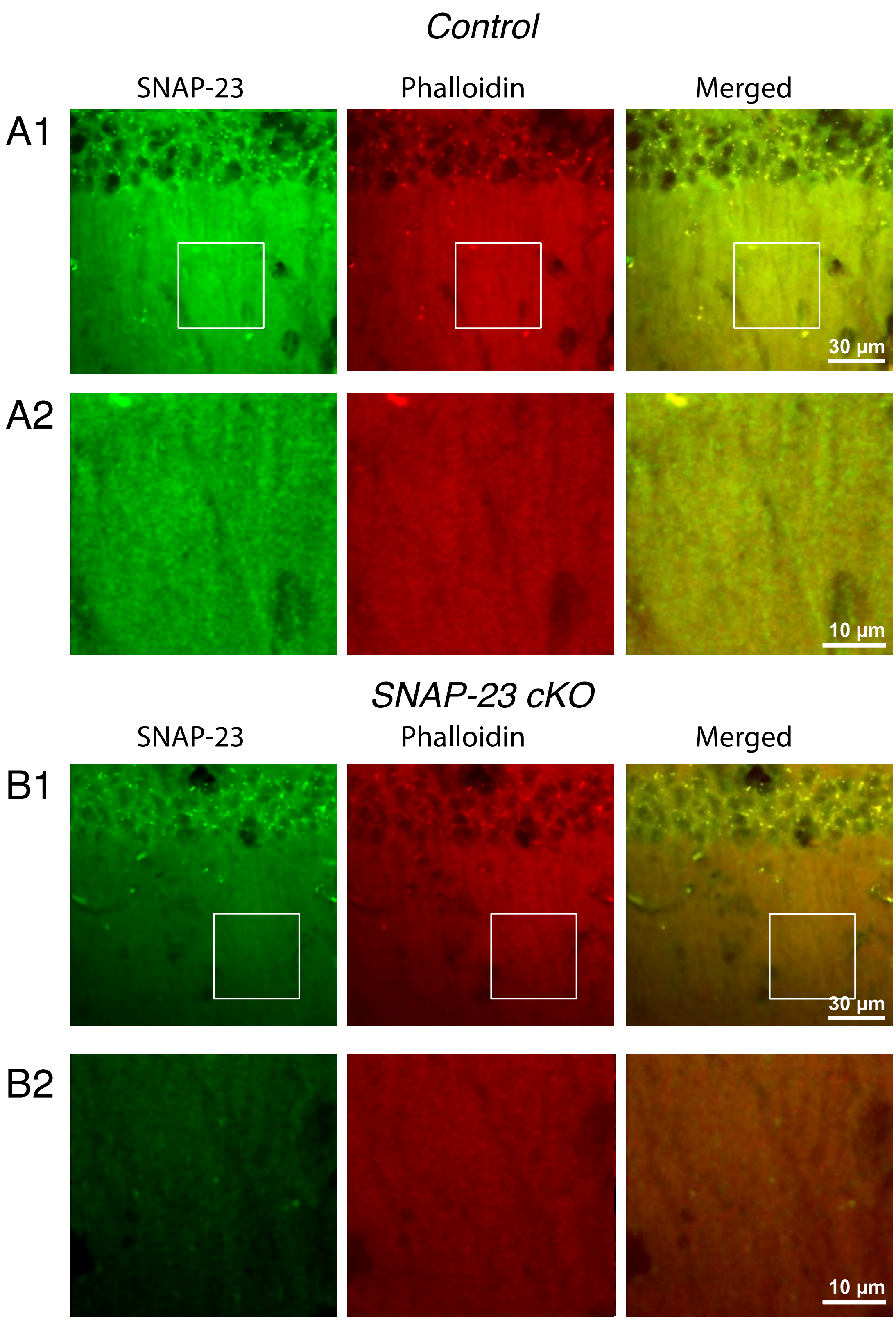


Supplemental Figure 1:

SNAP-23 is removed in CA1 cell distal dendrites. SNAP-23 (green) co-stain with rhodamine phalloidin (red) from control (A) and SNAP-23 cKO (B) brain sections.

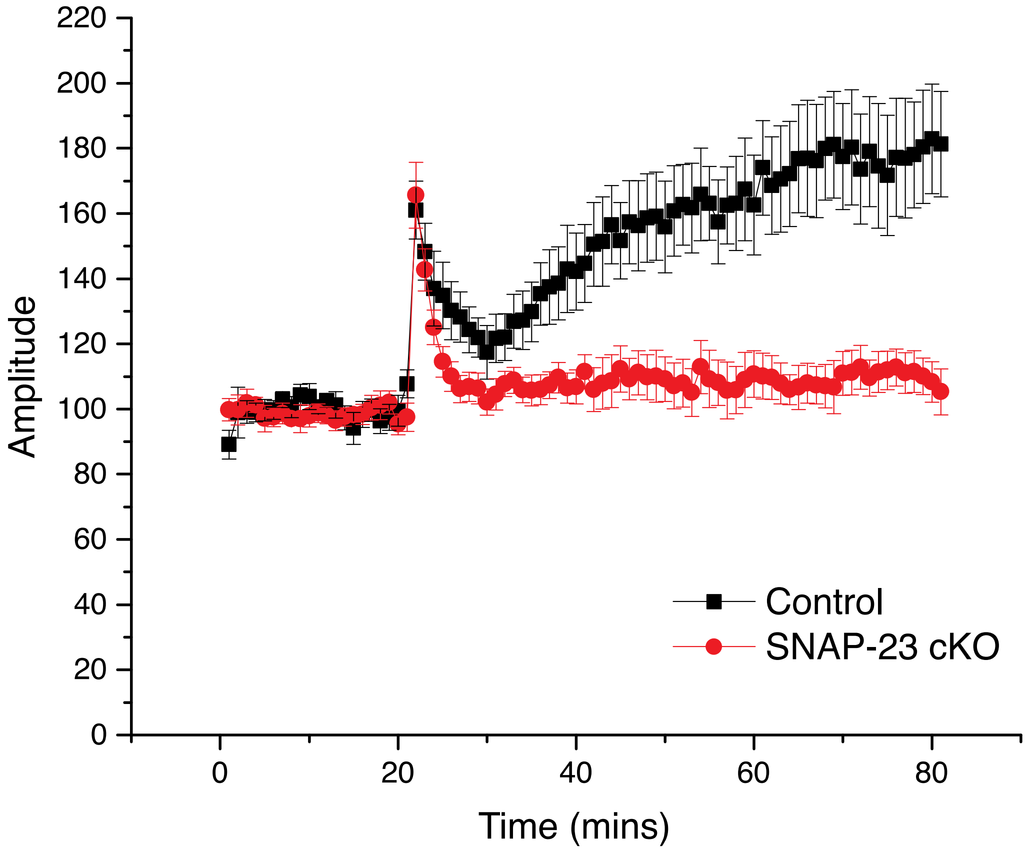


Supplemental Figure 2:

Average field EPSP amplitude normalized against baseline for SNAP-23 cKO and control.
